# Mountains as refugia: evidence from bumble bee brain transcriptomes

**DOI:** 10.1101/2022.09.09.507046

**Authors:** Kaleigh Fisher, Michelle A. Duennes, S. Hollis Woodard

## Abstract

As anthropogenic change continues to impact global biodiversity, the importance of rapidly identifying biodiversity refugia cannot be overstated. In this study, we employed a molecular test of the hypothesis that mountains serve as refugia for bumble bees against anthropogenic stressors. To explore this hypothesis, we compared stress-related patterns of gene expression in the brains of wild, pollen-foraging bumble bees of two species, *B. vosnesenskii* and *B. melanopygus*, collected at different elevations throughout the Sierra Nevada Mountain range in California, USA. We found evidence that the expression of several immune and detoxification genes is associated with elevational differences. This suggests that bees are experiencing differential exposure to stressors along an elevational gradient, which is an important criterion for identifying refugia across dynamic and heterogenous environments. This study thus provides evidence that mountains may serve as refugia for bumble bees in response to anthropogenic stressors, as has been detected for many other organisms.

## Introduction

Anthropogenic change, which includes climate change, disease spread, invasive species, and habitat loss due to urbanization, agricultural intensification, and deforestation, strongly impacts global biodiversity by disrupting species distribution patterns (Pimm et al., 2014; Isbell et al., 2017). In response to anthropogenic change, populations may decline (Barnosky et al., 2011; Ceballos et al., 2015), adapt to changing conditions (Johnson & Mushi-South, 2017; Gaynor et al., 2018), or shift to more hospital habitats (Chen et al., 2011). Refugia are defined as relatively stable places where species can withstand or adapt to disturbances and changing conditions (Gentili et al., 2015; Selwood & Zimmer, 2020). Although the concept of refugia has historically been used to describe behavioral responses to predation and species responses to glaciation events, it has also been used more recently in the context of anthropogenic change and biodiversity conservation. Specifically, it is hypothesized that relatively less-disturbed areas serve as refugia from anthropogenic stressors for many species (Morelli et al., 2020). Given rapidly increasing global change, these refugia are likely to become increasingly important locales for species persistence. Thus, identifying and preserving habitat types that serve as refugia is paramount for species persistence and conservation of biodiversity.

Mountains are habitats that can serve as refugia for many species in response to anthropogenic stress (Meng et al., 2019; Selwood & Zimmer, 2020; Huxley & Spasojevic, 2021). Although montane regions worldwide are being rapidly impacted by climate change (McCain & Colwell, 2011; Lenoir & Svenning, 2015; Rogora et al., 2018), they still appear to provide suitable habitats for many species relative to other habitat types (Elsen & Tingley, 2015; Guo et al., 2018; Elsen et al., 2020). This is because mountains are spatially heterogeneous, whereby microclimate and habitat features can vary substantially over short distances due to changes in elevation or slope aspects, which enable species to shift to some extent in response to extreme temperatures and reduced snowmelt (Elsen & Tingley, 2015; Meng et al. 2019). Moreover, they are relatively buffered from many other types of anthropogenic disturbances because montane ecosystems are often more difficult to develop for agriculture or urbanization, and in many areas they have stricter governmental protections (Nogués-Bravo et al., 2008; Elsen et al., 2018). For example, in the United States, nearly all subalpine or alpine areas are National Parks, which have strict regulations limiting land use and development (GAP Analysis Project, 2019). There are trade-offs related to living in high elevation areas, including a shorter growing season and lower oxygen levels (Jacobsen et al., 2020). However, for organisms that have evolved to withstand these ecological conditions (Liu et al., 2020), mountains can be relatively free of other stressors that are typically identified as key drivers of species decline (Mortiz et al., 2008; Leonir et al., 2008; Tingley et al., 2009).

Bumble bees (genus *Bombus*, family Apidae) are an excellent group to test whether montane areas can serve as refugia. An estimated 30% of the world’s ~250 bumble bee species are in decline (Arbetman et al. 2017), a pattern that is being strongly driven by anthropogenic stressors such as climate change and habitat loss (Goulson et al., 2015; Dicks et al., 2021). At the same time, many *Bombus* species have large habitat ranges that span both higher and lower elevation areas, allowing for within-species comparisons that consider the effects of montane gradients on environmental stress (Jackson et al. 2018, 2020). Bumble bees likely evolved in the Tibetan Plateau, then spread to many lower-elevation habitats around the world (William 1998; Williams et al. 2015). Today, they are found in both lowland and mountainous regions, but are particularly abundant and diverse in the latter (Williams, 1998; Fisher et al. 2021). This pattern is driven in part by their sensitivity to extreme heat, which is likely a holdover of their evolutionary history, and appears to be a major limiting factor of their distributions (Rasmont & Iserbyt 2012; Kerr et al. 2015; Soroye et al. 2020; Martinet et al. 2021). Over the last twenty years, several studies have demonstrated that many bumble bee populations are shifting to higher elevations and latitudes in response to climate change and habitat loss (Colla et al., 2006; Williams et al., 2009; Grixti et al., 2009; Kerr et al., 2015; Marshall et al., 2020; Suzui-Ohno et al., 2020). This pattern, which has been detected globally and across disparate species, strongly suggests that mountains are becoming increasingly important refuges for some bumble bee species.

Refugia can be challenging to identify because they are generally typified as having stable or positive changes to population densities or population genetics (Keppel et al., 2012), but these population responses can take time to reflect current anthropogenic stressors (Figueirido et al., 2019). Examining sublethal responses to stress using more dynamic molecular phenotypes, like gene expression, may help to overcome this challenge because these molecular changes can be detected over much shorter timescales, such as days or even minutes or seconds. Changes in the expression of nutritional, pathogenic, and immune canonical genetic pathways are all reflective of exposure to sublethal stressors in bees (Schmehl et al., 2014; Grozinger & Zayed, 2020; Costa et al., 2020; Costa et al., 2022), because these pathways play key roles in organizing biological processes, like detoxification, that allow bees to withstand these stressors (López-Osorio & Wurm 2020; Grozinger & Zayed, 2020). Moreover, more dynamic, organismal responses ultimately impact individual fitness and population level processes, including decline (Wikelski & Cooke 2006; Cooke et al. 2013). Analysis of gene expression, focusing on diagnostic biomarkers of stress, can thus be used to test for evidence of stress in wild populations (Grozinger & Zayed, 2020). Although experimental studies have begun to explore anthropogenic stressors, like pesticide application (Li et al., 2019; Gao et al., 2020; Costa et al., 2022) or thermal extremes (Pimsler et al., 2020), in lab settings, few studies have assessed how environmental factors impact gene expression in wild populations (but see Trego et al., 2019).

In this study, we employed a molecular test of the hypothesis that mountains serve as refugia for bumble bees against anthropogenic stressors. To explore this hypothesis, we compared stress-related patterns of gene expression in the brains of wild, pollen-foraging bumble bees of two species, *B. vosnesenskii* and *B. melanopygus*, collected at different elevations throughout the Sierra Nevada Mountain range in the Western United States. Most of the Sierra Nevada has federal or state protective status (Weixelman et al., 2011), with the highest-elevation areas all being National Park lands. However, the range is located between the Central Valley to the west, which is dominated by intensive agriculture, and the Great Basin to the east, which is desert habitat that is relatively uninhabitable for bumble bees (Thorp, 1983). Our two focal species are both widespread species that are found across a range of habitat types throughout the Western United States and are among the most abundant bumble bee species at various elevations throughout their range (Thorp, 1983; Koch et al., 2012; Strange & Tripodi, 2019; Cole et al., 2020; Fisher et al., 2021). Between the Central Valley and higher elevations in the Sierra Nevada, both species exist along a strong gradient of anthropogenic disturbance, with bees living at lower elevations exposed to greater anthropogenic stressors, like reduced resource quality, pesticide exposure, and extreme temperatures (Kremen et al., 2002; Williams et al., 2012). Given the challenges of studying gene expression in wild, free-living organisms (Alvarez et al. 2015), we used a relatively conservative methodology to maximize our ability to detect patterns of expression associated with clinal variation (Tito et al., 2020). We focused our study on the brain because gene expression in this tissue reflects stress-related processes like detoxification and immune response in insects (Costa et al., 2022) and because it has evolved mechanisms to maintain homeostasis (Treherne & Pichone, 1972). Thus, any changes in stress-related genes in the brain more likely reflect stressors encountered in their environment. We also focused specifically on expression of canonical immune and detoxification genes (Xu et al., 2013), which are well-annotated in the available bumble bee genomes (Sadd et al. 2015) and have the greatest amount of functional information to support their role in responding to stressors (Barribeau et al., 2015; Amezian et al., 2021). Based on the hypothesis that the Sierra Nevada Mountains are a refuge for bumble bees, we predicted that the expression of canonical, stress-related genes would differ across the elevational gradient sampled in our study, which would provide evidence that bumble bees are exposed to different levels or types of stress along elevation gradients. This type of evidence, whereby variation in response to stress across continuous, heterogeneous landscapes is demonstrated, is a critical criterion for identifying refugia in ecologically meaningful ways that can be used to support conservation policy (Keppel et al., 2012).

## Methods

### Bumble bee sample collection

In the Spring and Summer of 2017, we sampled a total of 78 workers from two bumble bee species that are common throughout the Sierra Nevada Mountains: *B. vosnesesnkii* (n = 45) and *B. melanopygus* (n = 33). These two species are the most abundant species in the state of California (Thorp, 1983; Fisher et al., 2021), are both members of the subgenus Pyrobombus, and shared a common ancestor < 20 million years ago (Hines 2008). We collected *B. vosnesenskii* workers from nine sites ranging in elevation from 366 m to 2675 m, and *B. melanopygus* workers from eleven sites ranging in elevation from 455 m to 2428 m (Figure 1). All individuals collected had pollen in their corbiculae, a sign that they were all actively foraging for pollen, in an attempt to minimize variation in brain transcription between pollen and non-pollen foragers. Bees were placed directly in liquid nitrogen and then stored at −80°C upon return from the field. We also recorded which plant species bees were collected from.

**Figure 1:**
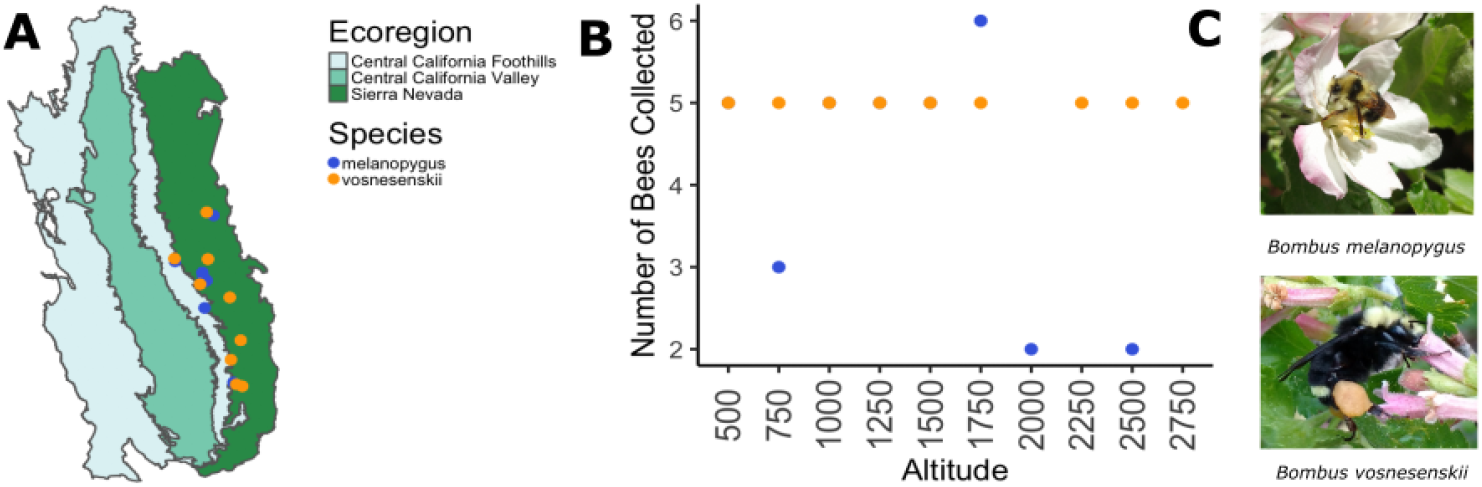
A) Map showing sites where bumble bee workers were collected based on USGS Level III Ecoregions. B) Number of individuals collected for each species at each altitude. C) Photographs of bumble bee species used in this study.

### Brain dissections, transcriptome sequencing, and assembly

We used whole brains for our RNAseq analysis to identify broad patterns of gene expression in this tissue type. Brains were dissected on dry ice using sterilized dissection tools cleaned with RNAsezap followed by 70% ethanol. Suboesophageal ganglia and glandular tissue surrounding the brain were removed. RNA was extracted using TRIzol and Qiagen RNeasy kits, with an RNase-free DNase treatment, following manufacturer’s instructions. Sample quality and integrity was determined by agarose gel electrophoresis, the Agilent 2100 bioanalyzer and nanodrop. RNA sequencing libraries for each brain (n = 78) were prepared with the following procedure: Samples were enriched using oligo(dT) beads, then fragmented randomly with fragmentation buffer. cDNA was then synthesized using the mRNA template and random hexamers primer, after which a custom second-strand synthesis buffer (Illumina), dNTPs, RNase H and DNA polymerase I were added to initiate second-strand synthesis. Finally, a series of terminal repair, A ligation and sequencing adaptor ligation were performed for final cDNA library construction. Quality control using Qubit 2.0, 2100 Agilent bioanalyzer and qPCR was performed, and then libraries were pooled and sequenced on an Illumina Novaseq Platform using 150 paired-end reads, which generated ~10 G raw reads for each sample. Data was then filtered to remove low quality reads (uncertain nucleotides > 10%; base quality < 20 constitute more than 50% or read), and adapter sequences. Trimmed and filtered reads were then aligned to the *B. impatiens* v2.0 reference genome using HISAT2.0 with splice sites and exon boundaries derived from the *B. impatiens* Official Gene Set (OGS) v1.0. Gene expression was quantified using featureCount v1.5.0-p3 (Liao et al., 2014) to determine the number of reads uniquely mapping to exons and summed at the gene level using gene features annotated in the *B. impatiens* OGSv1.0. Next, the number of fragments per kilobase of transcript per million mapped reads (FPKM) for each gene was calculated based on the length of the gene and reads count mapped to this gene. Library preparation, sequencing, and assembly were performed at Novogene Co. Ltd. (Sacramento, CA, USA).

### Data analysis

#### 1. Differential gene expression (DGE) analysis

We performed differential gene expression analyses on normalized transcript counts using the DESeq2 pipeline (Love et al., 2014) which uses median of ratios to normalize counts across samples according to sequencing depth and RNA composition. We considered the location workers collected from (site), the elevation where they were collected (elevation), and the plant species they were collected on (plant) as variables that could explain variation in gene expression between samples. We performed separate analyses for each species because we did not have species overlap at most sites. Moreover, when we analyzed species together, we found that species explained 49% of variation in gene expression after quality control analyses, which overshadowed any effects of environmental variables. We performed principle component analyses on rlog transformed counts to assess similarity between samples and visualize sources of variation in our dataset and to better inform model selection. For the *B. vosnesenskii* dataset, all three environmental variables were linear combinations; thus, we only included elevation in the DeSeq model (Love et al., 2014) because this was the environmental variable of interest. For the *B. melanopygus* dataset, location was a linear combination of plant and elevation, so only plant and elevation were included in the DeSeq model. We then used likelihood ratio tests between the full and reduced models to test whether elevation explained significant variation in gene expression, with a p-value < 0.05 being considered significant after FDR correction.

#### 2. Candidate gene enrichment

We generated a list of 218 candidate genes from published gene annotations in *B. impatiens* (Sadd et al., 2015), a *Bombus* species that is in the same subgenus (Pyrobombus) as *B. vosnesenskii* and *B. melanopygus* (Cameron et al., 2007). Our candidate gene list was comprised of 184 immune-related genes (Barribeau et al., 2015) and 24 detoxification genes, which included twelve genes in the Glutathione-S-Transferase (GST) family (Sadd et al., 2015) and 22 genes in the CYP3 p450 (Darragh et al., 2021; Haas et al., 2022).

To increase the robustness of our results, we performed differential gene expression (hereafter DGE) analysis using elevation as a continuous variable, rather than binning elevation into discrete units, which resulted in generating fewer differentially expressed genes. Similarly, we only included genes with existing evidence of being related to a stress-related function based on genome annotations from the *Bombus impatiens* (Sadd et al., 2014; Barribeau et al., 2015). Due to these strict criteria, we likely underestimated the total number of genes that are associated with elevation that may be stress-related. For example, we did not include heat shock proteins in our analysis because there are not comprehensive gene annotations for this family currently available, despite their known role in thermal response in bumble bees (Kim et al., 2008; Pimsler et al., 2021). For greater detail on how we generated gene lists, please refer to Appendix 1. To test whether the immune or detoxification gene families were enriched for either *Bombus species*, we generated contingency tables followed by Fisher’s exact tests.

#### 3. Candidate gene patterns

We visualized candidate gene patterns using principle component analyses by considering all candidate genes, regardless of whether they were differentially expressed based on likelihood ratio tests. We then performed Wald tests to estimate how gene expression varied with changes in elevation along a continuous elevational gradient, with each log2 fold change in gene expression representing one unit of change in elevation.

## Results

We analyzed the expression of 12,925 genes across 78 individual bumble bee worker brains, which represents 98% of the genes in the *B. impatiens* genome (Sadd et al. 2015). Using likelihood ratio tests, we found 251 genes upregulated and 284 downregulated in *B. vosnesenskii*, for a total of 535 genes differentially expressed in response to changes in elevation. We found 141 genes upregulated and 176 downregulated in *B. melanopygus* (Table 1).

**Table 1:**
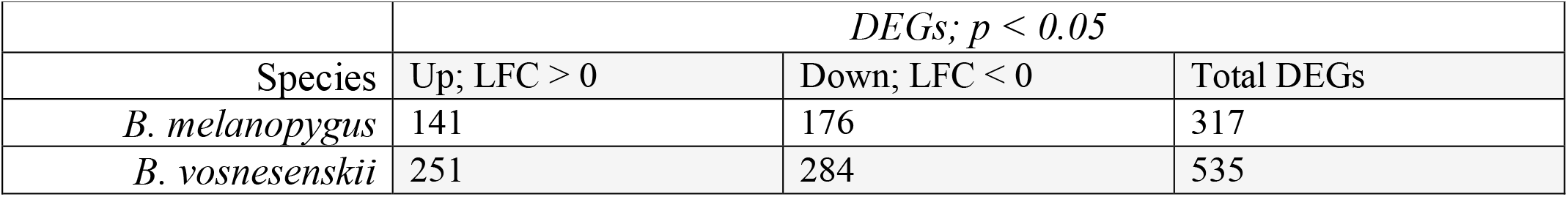
Number of genes that were differentially expressed *Likelihood ratio test; p-value < 0.05*

### Immunity and detoxification gene patterns

We did not detect general patterns of gene expression based on all immunity and detoxification genes in response to elevation for *B. vosnesenskii* along the first principle component axis. There was some variation along the second axis in response to elevation, but several outliers at 750, 2250 and 2500 m may have obscured any conclusive patterns (Figure 2). We found more pronounced differences along PC1 in *B. melanopygus* based on all immunity and detoxification genes, but not PC2. Gene expression profiles were more similar in bees collected from the highest and lowest elevations compared to gene expression profiles of bees collected in more median elevations (1250-1750 m) (Figure 3).

**Figure 2:**
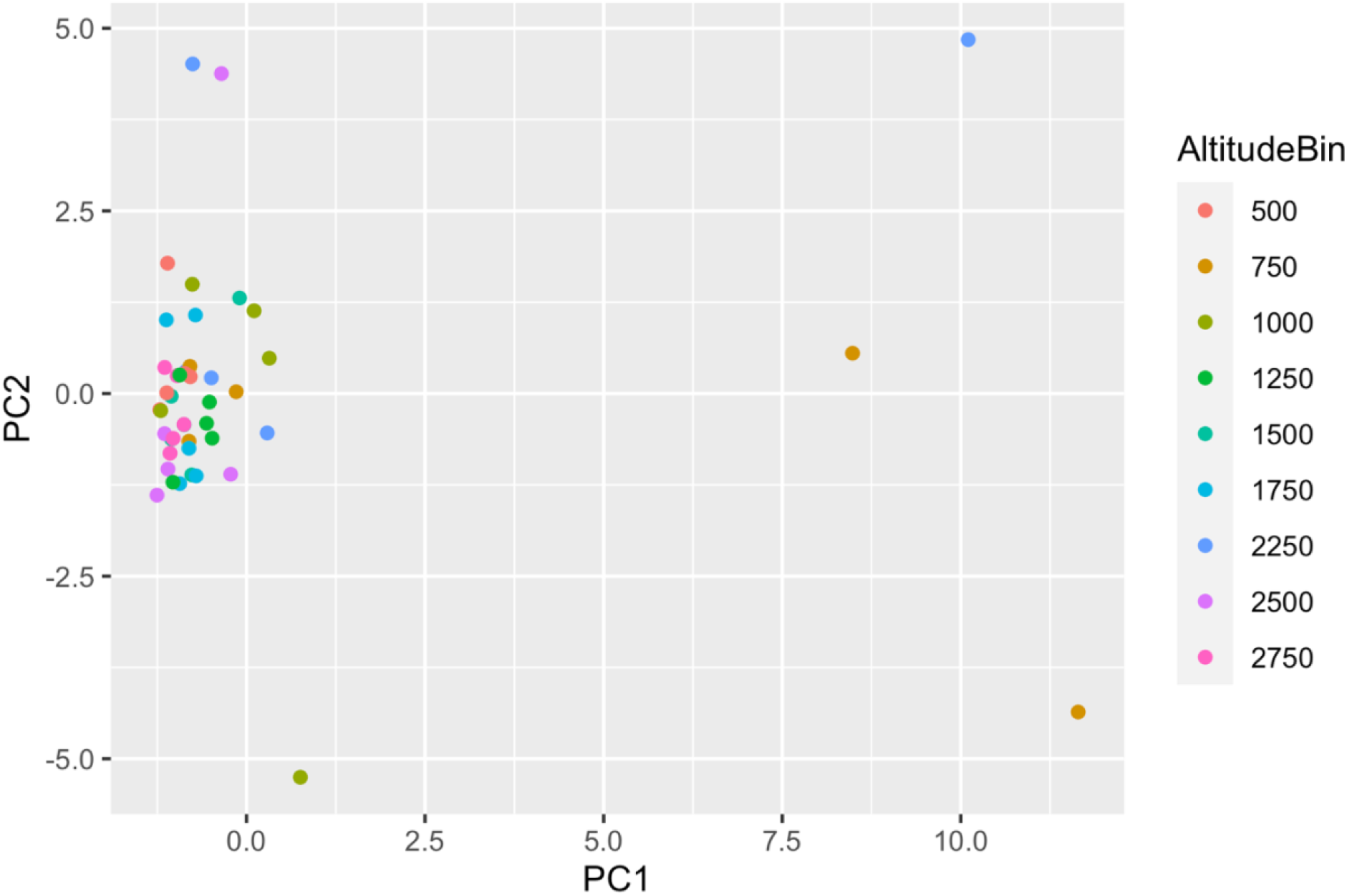
PCA showing similarity between *B. vosnesenskii* samples based on normalized counts of 207 candidate genes (Immune: 175 genes; Detoxification: 32 genes). Note: Elevation was binned into 250 m bins for visual clarity.

**Figure 3:**
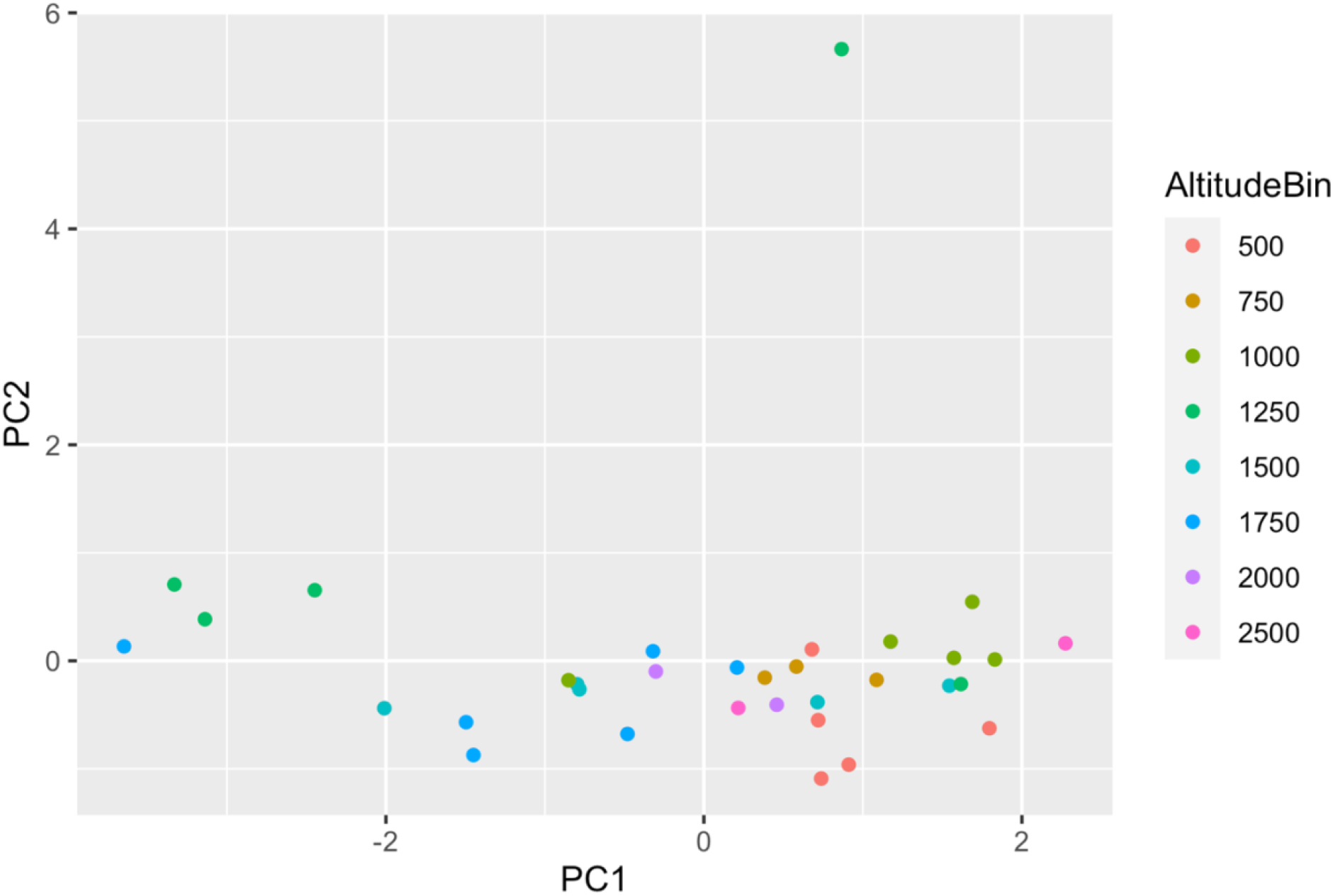
PCA showing similarity between *B. melanopygus* samples based on normalized counts of 205 candidate genes (Immune: 173 genes; Detoxification: 32 genes). Note: Elevation was binned into 250 m bins for visual clarity.

### Immunity and detoxification gene representation

We detected expression of 213 of the 218 genes in our candidate gene lists, which comprised 179 immune genes and 34 detoxification genes. In the *B. vosnesenskii* dataset, we detected expression of 170 immune genes and 30 detoxification genes. Nine immune genes and two detoxification genes were found to be differentially expressed in response to changes in elevation. In the *B. melanopygus* dataset, we detected expression of 168 immune genes and 30 detoxification genes. Five immune genes and two detoxification genes were found to be differentially expressed relative to changes in elevation. Neither of these gene functional categories were found to be overrepresented in the dataset for either species (Table 2). The expression of about half of the differentially expressed immune genes were positively correlated with elevation in both species (Table 3). Only one of the differentially expressed genes, CLIPD-like gene, was detected as being differentially expressed in association with elevation in both species, but it had contrasting expression patterns in response to elevation. However, there were several genes that were differentially expressed in both species that have putatively similar functions. For example, we found differential expression of genes in the relish gene family and RNA helicases in both species; expression of these genes was positively correlated with elevation. Both species also had IAP (inhibitor of apoptosis) genes, though they also exhibited contrasting patterns in response to elevation between the two species. There were two p450s differentially expressed for each species. Both the p450s were negatively correlated with elevation in *B. melanopygus* whereas one was negatively correlated, and one was positively correlated with elevation in *B. vosnesenskii*.

**Table 2:** Over-representation analysis for gene categories of interest

| Species | *Immune* | *Detoxification* |
| --- | --- | --- |
| *B. melanopygus* | **5** (168)<br>*P* = 0.50 | **2** (30)<br>*P* = 0.21 |
| *B. vosnesenskii* | **9** (170)<br>*P* = 0.44 | **2** (30)<br>*P* = 0.44 |

Bold number represents number of DEGs in that category, parentheses indicate total genes of that category found in the model \* Indicates significance for Fisher's Exact Test ( $p < 0.05$ )

**Table 3:** List of differentially expressed genes in each species in response to changes in elevation

| Species | Gene Category | Gene Name | Log2Fold Change | p-adj. | Gene ID |
| --- | --- | --- | --- | --- | --- |
| *B. melanopygus* | Immune | REL1A* | + | 0.00277 | LOC100740410 |
|  | Immune | CLIPD-like* | - | 0.01604 | LOC100741694 |
|  | Immune | RM62I* | + | 0.00005 | LOC100746451 |
|  | Immune | TPX2 | + | 0.02440 | LOC100747832 |
|  | Immune | IAP5* | - | 0.00229 | LOC100748174 |
|  | p450 | probable cytochrome P450 6a14* | - | 0.00003 | LOC100740683 |
|  | p450 | cytochrome P450 9e2 | + | 0.03013 | LOC100749156 |
| *B. vosnesenskii* | Immune | IAP2* | + | 0.03726 | LOC100740635 |
|  | Immune | CLIPD-like* | + | 0.04929 | LOC100741694 |
|  | Immune | REL1B* | + | 0.04413 | LOC100741932 |
|  | Immune | RM62H* | + | 0.04315 | LOC100742333 |
|  | Immune | CTLGA1 | - | 0.00226 | LOC100744252 |
|  | Immune | DEF1 | + | 0.00453 | LOC100744436 |
|  | Immune | PASHA | - | 0.00563 | LOC100745100 |
|  | Immune | SPZ1B | - | 0.04803 | LOC100747254 |
|  | Immune | APG4B | - | 0.00903 | LOC100748069 |
|  | p450 | probable cytochrome P450 6a14* | + | 0.00060 | LOC100740804 |
|  | p450 | cytochrome P450 6a2-like | - | 0.00604 | LOC100743789 |

## Discussion

There is growing evidence that montane ecosystems can serve as a refugia for many species. Biodiversity refugia are frequently identified through species monitoring, whereby habitats that support stable or growing population numbers are considered refugia (Keppel et al., 2012; Selwood & Zimmer, 2020). However, refugia are complex in that they are not discrete spatial homogenous units, but support species to varying degrees over continuous, heterogeneous landscapes (Keppel et al., 2012). Moreover, the locations that serve as refugia may change dynamically through time, and may become increasingly important with ongoing climate change, as is predicted for mountains. Here, we performed one of the first studies examining the relationship between environmental variation and gene expression in wild bees (but see Brunner et al., 2013; Tsvetkov et al., 2021), to test the hypothesis that mountains serve as refugia for bumble bees against anthropogenic stressors. We show that in two species of bumble bees living in the Sierra Nevada Mountains and adjacent foothills, the expression patterns of several canonical stress-related genes are associated with changes in elevation. This suggests that bees are experiencing differential exposure to stressors along an elevational gradient, which is an important criterion for identifying refugia across dynamic and heterogenous environments (Keppel et al., 2012). This study thus provides evidence that mountains may serve as a refugia for bumble bees in response to anthropogenic stress, as has been detected for many other organisms (Chen et al., 2011; Selwood & Zimmer, 2020). This supports the broader argument that montane habitats are particularly important landscapes to target in land-based conservation (Kollmair et al. 2005).

Bumble bees are currently facing numerous anthropogenic stressors, including a lack of nutrient-rich food resources, exposure to pesticides and pathogens, and climate change (Goulson et al., 2015). The sublethal effects of these stressors can be detected via changes in gene expression patterns in controlled laboratory studies (Tsvetkov et al., 2021; Costa et al., 2022). Although there is evidence that higher elevations and latitudes may buffer bumble bee populations against all of these anthropogenic stressors (Jackson et al., 2018; Pimsler et al., 2020), we do not have conclusive gene candidates for response to lack of food resources and climate change, in contrast to the strong support that suggests pesticide and pathogen exposure impact detoxification and immune genetic pathways, respectively (Barribeau et al., 2015; Troczka et al., 2019; Haas et al., 2022). There are known challenges associated with analyzing gene expression in wild animals, including high levels of unexplained variation that make functional inferences more difficult (Alvarez et al. 2015). To account for this in our study, we set relatively stringent criteria whereby we only considered genes that were previously identified from the *B. impatiens* genome to likely be involved in stress-related processes. Moreover, we analyzed differential gene expression in response to elevation as a continuous variable. Thus, our results only reflect genes that vary across the entire elevation gradient, making our results more robust.

We found evidence for changes in gene expression of stress-related genes in response to elevation in two focal bumble bee species, *B. melanopygus* and *B. vosnesenskii*. When we considered all stress-related genes *B. melanopygus*, we found that bees collected at intermediate elevations (1,250-1,750 m) had greater similarity in overall expression profiles, whereas bees collected from lower and higher elevations were more similar, compared to those of intermediate elevations. There were not such clear patterns in overall expression profiles for *B. vosnesenskii*. These contrasting patterns between the two species may have several non-mutually exclusive explanations. First, species-specific differences in foraging phenology may contribute to the patterns observed. *B. melanopygus* workers begin foraging earlier in the season than *B. vosnesenskii* (Williams et al. 2014), and subsequently visit typically different plants (Cole et al., 2020). As a result, we were only able to collect both species from two out of eighteen total sites, most of which had different floral resources available. Floral resource quality and geographic location likely also impact expression of stress-related genes (Mao et al., 2013; Vannette et al., 2015), but we were unable to disentangle these factors from elevation because of the collinearity of these variables with elevation in our study. Second, homologous genes that are shared between two species may have similar functions, such as being involved in immune response, but may operate and be regulated in slightly different ways. Thus, stress-response genes shared by the two species considered in this study may display different expression patterns despite sharing the same basic function. Despite these limitations, our ability to detect evidence that several key stress-related genes are differentially expressed in both species in response to elevation suggests that they may contribute to bumble bees’ responses to anthropogenic stressors and should be further investigated.

We focused this study on immune and detoxification genes because of their known functions in insect response to pesticide and pathogen exposure (Barribeau et al., 2015; Troczka et al., 2019; Haas et al., 2022). The bumble bee genome contains all of the components of the typical immune pathways in insects (Sadd et al., 2015; Barribeau et al., 2015), though it contains reduced representation of genes amongst the immune pathways, as well as a reduced repertoire of detoxification genes compared to many other insects (Sadd et al., 2015). All of these genes have conserved functions related to either detoxification or immune response, though the specific mechanisms and gene expression patterns of each gene are not completely resolved. Most of what we know about the functions of these genes is from research on the fruit fly (*Drosophila*) and the Western honey bee (*Apis mellifera*), though some gene functions have been directly characterized in bumble bees (Haas et al. 2022). Changes in gene expression may reflect a negative consequence of the stressor on physiological processes, or the activation of a physiological pathway that enhances resilience to a stressor (Vanette et al., 2015; Grozinger & Zayed, 2020). Thus, upregulation of stress-related genes in the brains of bees collected at higher elevations may be related to an increased resilience against pesticides or pathogens. Alternatively, upregulation in response to stressors may indicate that the bees are physiologically stressed and as a result they may ultimately suffer fitness consequences. Importantly, we cannot differentiate between these scenarios and more functional studies of stress-related genes are necessary to do so.

Two immune genes, REL1B and SPZ1B, which we found differentially expressed in our study, were also differentially expressed in another study that looked at pesticide exposure and pollen diet quality on gene expression in bumble bee brains (Costa et al., 2022). REL1B was downregulated in response to pesticide exposure and pollen quality but upregulated in *B. vosnesenskii* brains in our study in response to higher elevations. SPZ1 in contrast was upregulated in response to pollen quality in Costa et al. (2022) and downregulated in response to higher elevations in our study. Two p450 detoxification genes were also differentially expressed in both our study and Costa et al., 2022. p450s are involved in pesticide detoxification in insects (Amezian et al., 2021). One p450 gene was upregulated in Costa et al. (2022) in response to pesticide exposure and downregulated in *B. melanopygus* workers collected at higher elevations whereas the other p450 was upregulated in response to pesticide exposure in Costa et al. (2022) and upregulated in *B. vosnesenskii* at higher elevations. Given that pesticide exposure decreases with increasing elevations in the Sierra Nevada, these results may suggest that bees collected at higher elevations were more buffered from pesticides, and that these two p450 genes may serve as indicators of this trend. Anthropogenic stressors can have combinatorial effects in bumble bees (Costa et al., 2022), and thus impact molecular pathways in interactive ways. Though bees at higher elevations in the Sierra Nevada likely experience fewer anthropogenic stressors, more experimental work is needed to disentangle these patterns. Given that the immune genes REL1B and SPZ1, and the p450 genes, were detected in both of these studies, they are all excellent candidates for future experimental work to help understand what pathways are involved in anthropogenic stress response in bees. Subsequently, they may also serve as important biomarkers for stress response in bumble bees (Grozinger & Zayed, 2020). Further research that characterizes the precise function of these candidate genes will help to contextualize the biological significance of why there are differences in expression across various elevations.

As the planet continues to become less hospitable for many organisms, the importance of identifying and protecting habitats that serve as refugia for biodiversity cannot be overstated. Standard approaches for identifying refugia involve integrating data on habitat characteristics (e.g., resource availability, climatic conditions) that support species persistence with species distribution data over relevant spatial and temporal scales (Keppel et al., 2012, Morelli et al., 2020). Dynamic molecular markers like changes in gene expression complement this framework because they can reflect the health of populations long before population declines are detected (Tilman et al., 1994). This study provides evidence that gene expression can also be used to look for evidence of refugia, which is a complementary approach to more standard methods. Although it can be difficult to interpret gene expression in wild organisms because it is impossible to control many of the interacting factors that influence gene expression, our study demonstrates that it is possible to link patterns of expression for well-characterized genes with environmental variation across heterogeneous landscapes (Trego et al., 2019). Thus, dynamic molecular phenotypes, like gene expression, can be used as an additional conservation tool for identifying and justifying the protection of places that serve as refugia for species.

## Appendix S1

The candidate gene list was generated by combining information on gene annotations from the following sources: immune gene annotations were provided in Sadd et al., 2014. CYP3 p450 genes annotations were obtained from Darragh et al., 2021 and were then searched for in the gene database on NCBI for *Bombus impatiens* (https://www.ncbi.nlm.nih.gov/data-hub/gene/table/taxon/132113/). GSTs were also searched for in the NCBI gene database for *Bombus impatiens*.

